# Poor adult nutrition impairs learning and memory in a parasitoid wasp

**DOI:** 10.1101/2021.01.09.426073

**Authors:** Hossein Kishani Farahani, Yasaman Moghadassi, Jean-Sebastien Pierre, Stéphane Kraus, Mathieu Lihoreau

## Abstract

Animals have evolved cognitive abilities whose impairment can incur dramatic fitness costs. While malnutrition is known to impact brain development and cognitive functions in vertebrates, little is known in insects, whose small brain appears particularly vulnerable to environmental stressors. Here, we investigated the influence of diet quality on learning and memory in the parasitoid wasp *Venturia canescens*. Newly emerged adults were exposed for 24h to either honey, sucrose solution 20%, sucrose solution 10%, or no food, before being conditioned in an olfactory associative learning task in which an odor (orange) was associated to a reward (host larvae). Wasps fed honey showed 3.5 times higher learning performances and 1.5 times longer memory retention times than wasps fed sucrose solutions and starved wasps. Poor diets also reduced longevity and fecundity. Our results demonstrate the importance of early adult nutrition for optimal cognitive function in these parasitoid wasps that must quickly develop olfactory memories for choosing high quality hosts for their progeny.

## Introduction

Animals rely on various forms of learning and memories to exploit resources in their environment and adapt to changing conditions [1, 2, 3]. These cognitive abilities are sustained by brains that require large amounts of proteins to grow [4], but also lipids and carbohydrates for maintenance [5, 6, 7]. The process of learning, itself, imposes important energetic costs [8], and the formation of persistent (long-term) memories involves protein synthesis [9, 10]. Therefore, the ability of animals to acquire key nutrients in food is expected to directly impact their cognitive performances [11].

Malnutrition is known to affect cognitive functions in vertebrates (e.g. pigeons: [12], mice: [13], cats and dogs: [14]). In humans, for instance, high fat and caloric diets have been associated with hippocampal-dependent memory loss [15, 16]. By contrast, little is known about the cognitive effects of malnutrition in invertebrates. Insects, in particular, rely on an impressive range of learning and memory forms to interact socially and forage, and these cognitive abilities are implemented by only few neurons [17, 18]. The miniature brain of insects is thus particularly vulnerable to a range of environmental stressors, including poor nutrition [19]. For instance, in the fruit fly *Drosophila melanogaster*, larvae fed diets with unbalanced protein to carbohydrate ratios showed reduced learning performances in an aversive olfactory differential learning task [20]. In the Western honey bee *Apis mellifera*, adults fed pollen with a deficit in specific fatty acids (i.e. Omega-3, Omega-6) had impaired learning and memory performances in an appetitive olfactory differential learning task [21, 22]. These cognitive effects of malnutrition may incur particularly strong fitness costs in many solitary species where adults rely on learning and memory to find food and nourish their progeny by themselves.

Solitary parasitoid wasps, such as *Venturia canescens*, learn to associate an odor with a high-quality host to select nutritionally rich environments for the development of their offspring [23, 24, 25]. Before engaging in oviposition, adults encompass a critical period soon after emergence, where they need to find food [26, 27, 28]. Females typically acquire carbohydrates from nectar and honeydew [29, 30, 31]. Other key nutrients such as proteins, minerals and fat, are occasionally obtained from pollen [32]. Since olfactory memory formation is nutritionally demanding, we hypothesized that wasps fed highest quality diets would show the best cognitive performances.

Here, we tested whether diet quality during early adulthood affect the cognitive performances of *V. canescens* wasps. We experimentally exposed emerging females to one of four nutritional conditions of decreasing quality in terms of nutrient concentration, nutrient diversity, and energy content (honey – see composition in Table 1, sucrose 20%, sucrose 10%, no food). We then tested the impact of diet on cognition in a conditioning assay in which wasps had to associate an odor to a reward (host) in a flight tunnel. We further tested the influence of diet on fitness by monitoring the longevity and the reproductive success of these wasps.

**Table 1.** Honey composition. Honey analysis was made by ASA Laboratory (Tehran, Iran) based on ISIRI-7610, ISIRI-92 and European Honey Directive and the Codex Alimentarius Standard for Honey standards.

| Ingredients |  | Per 100 g |
| --- | --- | --- |
| Protein |  | 1.5 g |
| Carbohydrates |  | 70 g |
| Sugars | Glucose | 30 g |
|  | Fructose | 38 g |
| Fat |  | 0 |
| Fiber |  | 1 g |
| Vitamins | B6 | 4% |
|  | C | 3% |
|  | Riboflavin | 8% |
|  | Folate | 3% |
| Potassium |  | 50mg |
| Sodium |  | 10mg |
| Total Energy |  | 284 cal/mg |

## Results

### Wasps did not show innate odor preference

We first tested the influence of the nutritional condition on innate attraction to odors, by giving individual wasps a simultaneous choice between two odor sources (orange and vanilla) in a flight tunnel with two decisions chambers for 15 min (see details in Figure 1). The wasps did not display any preference for either odor, irrespective of their nutritional condition (χ_*2*_ ^2^ = 0.13, *P* = 0.93, N = 50). The proportion of wasps that made no choice (i.e. when the wasps did not fly after 5 mins in the tunnel) remained stable (30±2%) and was similar across nutritional conditions (χ_*2*_ ^2^ = 0.9, *P* = 0.69, N = 50). This indicates that the nutritional condition did not affect the motivation nor the motor activity of wasps in response to odorants. We therefore arbitrarily chose the orange odor as the conditioned odor (CS+) and the vanilla odor as the new odor (NOd) in all the subsequent behavioral tests.

**Figure 1.**
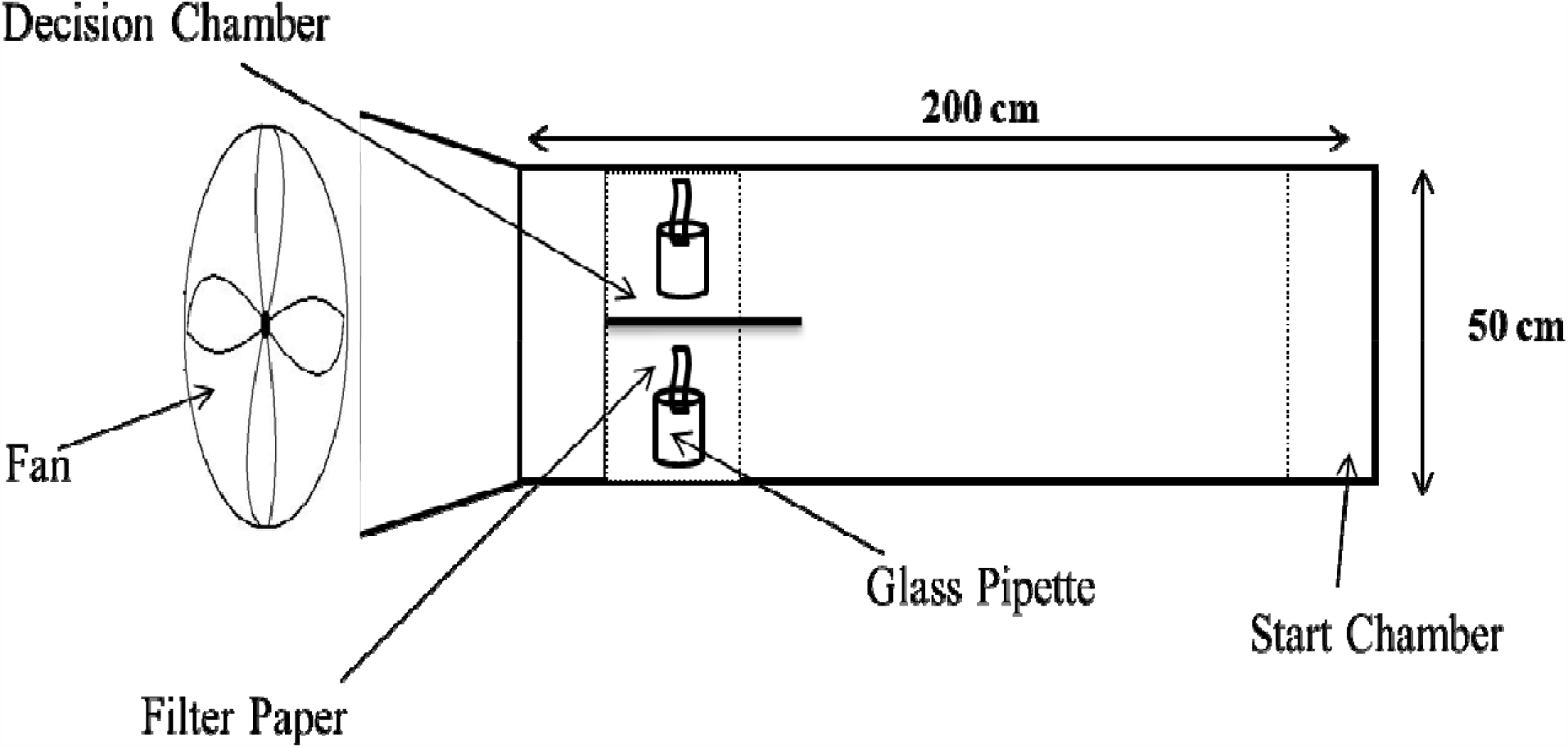
**Schematic view of the flight tunnel (top view)**. Individual wasps were introduced in the start chamber and observed choosing between the two odors displayed on filter papers in the decision chambers for 15 minutes.

### Wasps fed honey showed highest learning performance

We then tested the effect of the nutritional condition on olfactory learning. To do so, we first conditioned each wasp in the presence of 30 host larvae and the orange odor (CS+) for 2h. We then tested the conditioned wasps for odor preference by giving them a choice between the CS (orange) and the NOd (vanilla) in the flight tunnel for 15 min. Learning was observed in wasps exposed to each of the four nutritional conditions but at different magnitudes (Figure 2; Binomial GLM Feeding; χ ^2^=87.33, P<.0001; Conditioning: χ ^2^=14.48, P=0.0001; Feeding× conditioning: χ^2^=7.7, P=0.005). The highest proportion of correct choices for the CS (85±3%, N =50) was observed in wasps fed honey. This proportion decreased with diet quality, reaching intermediate levels in wasps fed sucrose diets (72±2%, N= 50), and a minimum level in starved wasps (62±2%, N = 50). The proportion of wasps that did not make a choice remained low across nutritional conditions (12 ± 2%, N=50; Figure 2), but increased with decreasing diet quality, reaching a maximum in starved wasps (29±3%, N =50; Binomial GLM Feeding; χ ^2^=12.3, P=0.006; Conditioning: χ^2^=8.93, P=0.002; Feeding× conditioning: χ^2^=7.61, P=0.005; Figure 2). Therefore, wasps fed highest diet quality showed highest learning performances.

**Figure 2.**
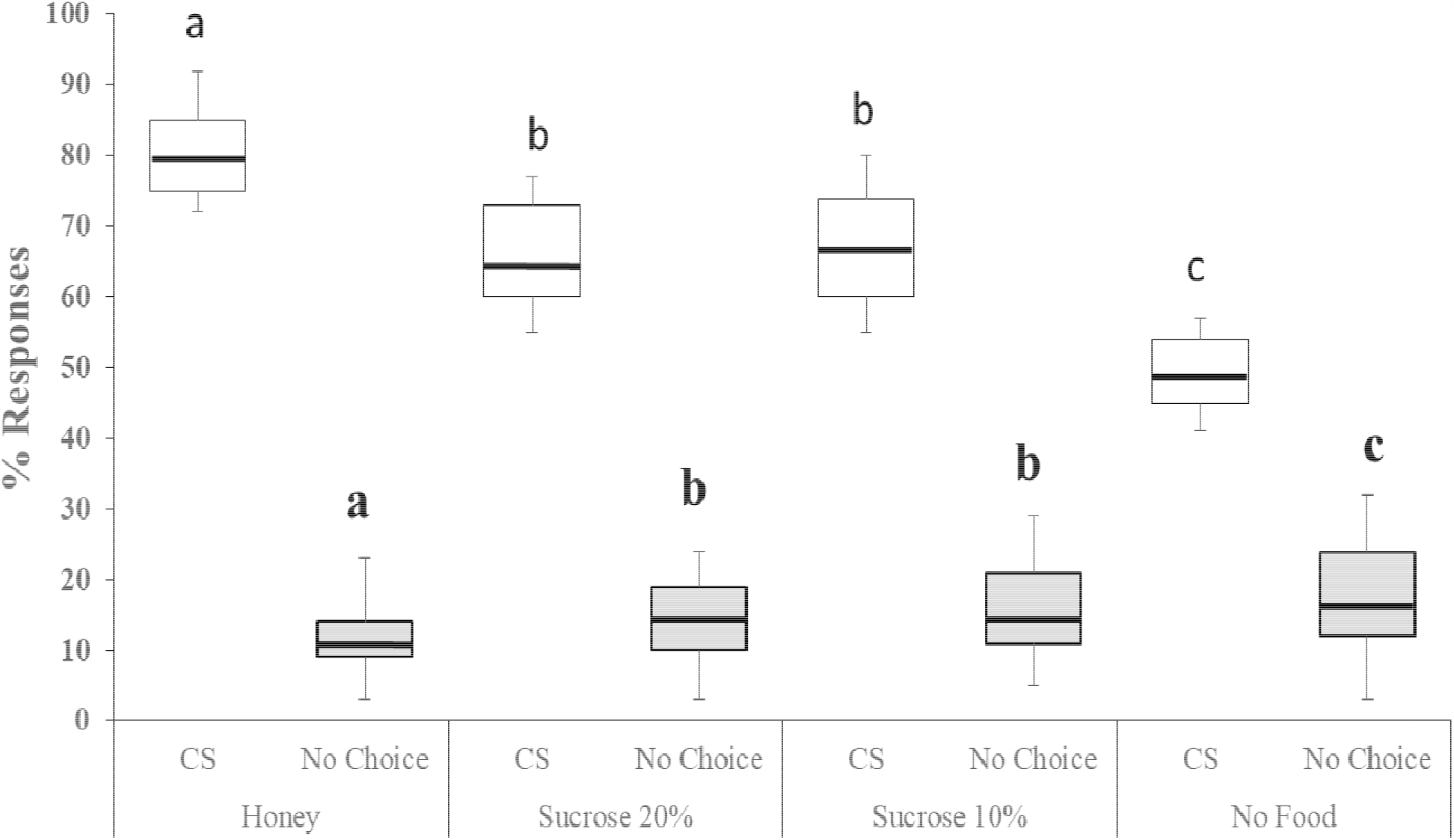
**Learning performances**. Percentages of correct choices for the conditioned stimulus (CS, orange odor) and no choices for each nutritional condition. Box plots represent the median (bold line), the interquartile range (length of box), and extreme, maximum and minimum, data points. Generalized Linear Models (GLM) were implemented with the binomial family error and logit link. Different letters above bar plots indicate significant differences between the treatments after Bonferroni correction (P = 0.0125). N = 50 wasps per nutritional condition (200 wasps in total).

### Wasps fed honey showed longest memory retention

We further tested the effect of the nutritional condition on olfactory memory retention by testing wasps in the flight tunnel at different time periods, between 2h and 30h post conditioning (Figure 3). Memory retention was significantly longer in wasps fed honey (27±3h, N= 600) than in wasps fed 20% (12±3h, N= 600) and 10% (17±2h, N= 600) sucrose solutions, and starved wasps (8±2h, N= 600; ANOVA: F= 302.2, P< 0.0001). Therefore, wasps fed highest diet quality showed longest memory retention.

**Figure 3.**
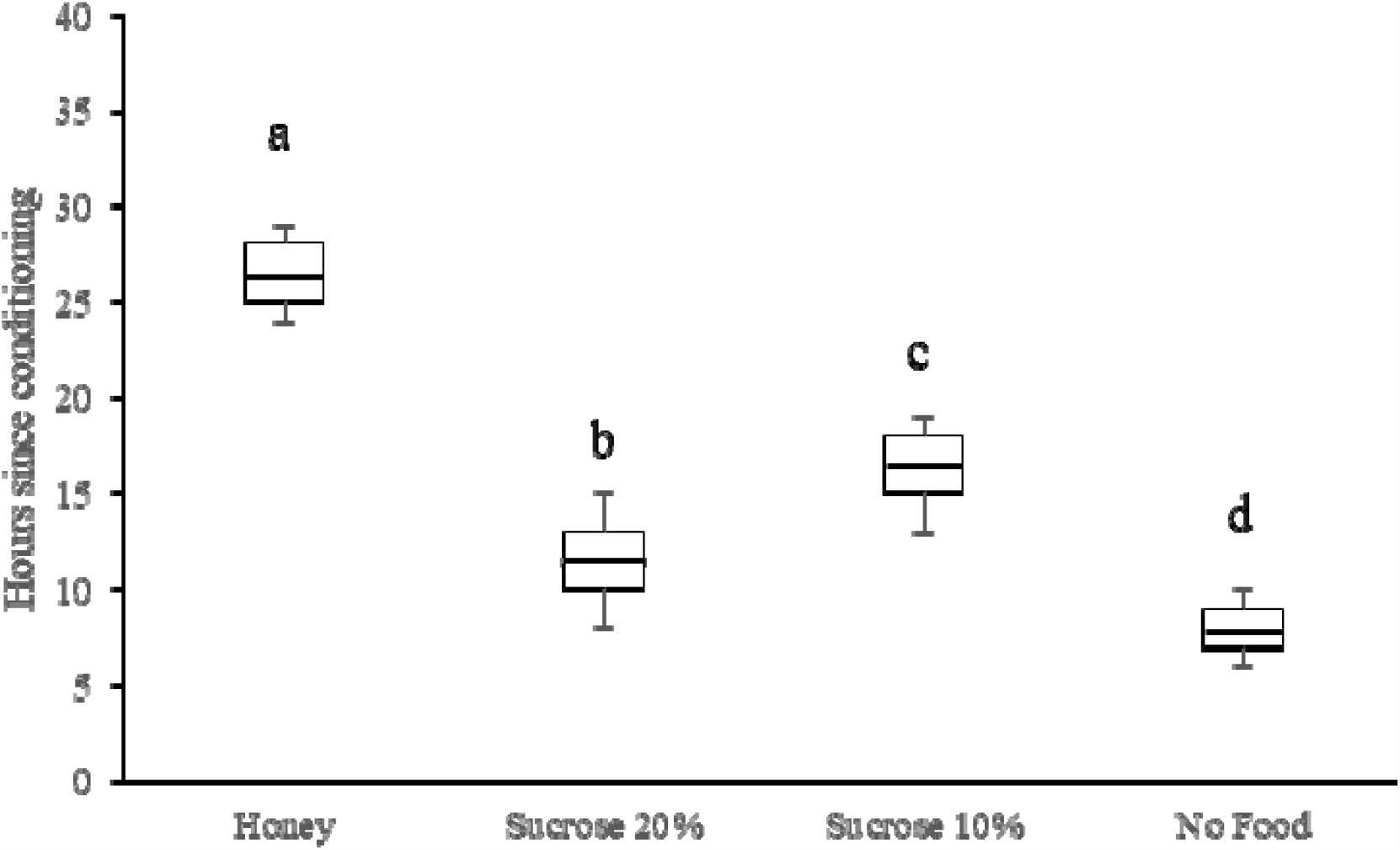
**Memory retention times**. Box plots show the median (bold line), the interquartile range (length of box), and extreme, maximum and minimum, data points. Memory retentions were compared using one way analysis of variance (ANOVA). Different letters indicate significant differences between the treatments after Bonferroni correction (P = 0.0125). N = 50 wasps per nutritional condition and time interval (2400 wasps in total).

**Figure 4.**
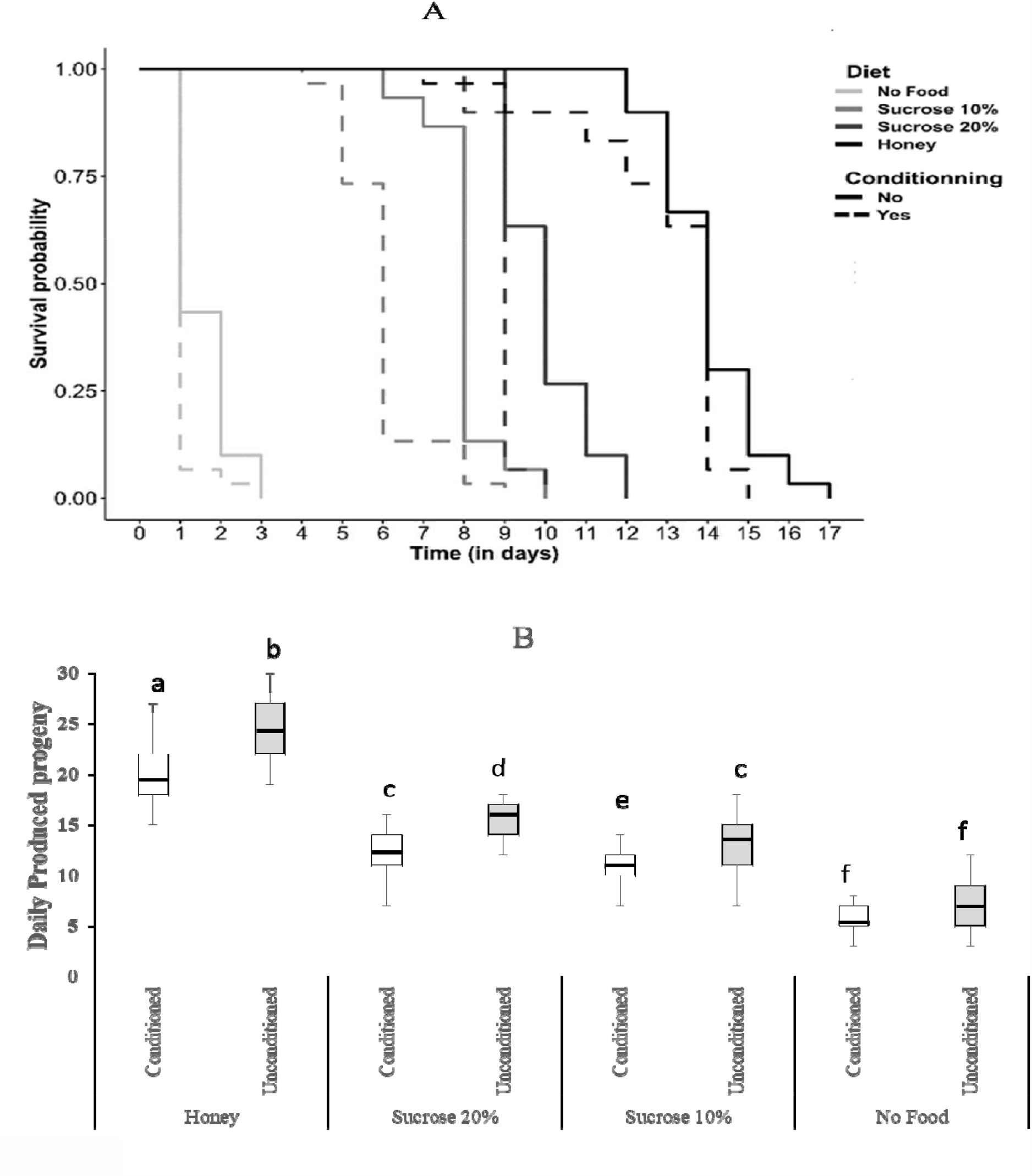
**Longevity and fecundity**. Effect of the nutritional condition (colors) and conditioning (solid or dashed lines) on survival probability (**a**) and fecundity (**b**) of female wasps. Survival curves were obtained from Kaplan Meier model (function survfit in R package “survival” (Therneau 2015)). In the boxplot, the central line is the median, the edges of the box are the 25th and 75th percentiles, the whiskers extend to the most extreme data points not considered outliers, outliers are represented by points. Median lifespan is indicated for each group in the fecundity graph. Different letters indicate significant differences between the treatments after Tukey post-hoc.

### Wasps fed honey had highest longevity and fecundity

We finally tested the effects of the nutritional condition (i.e. diets) and conditioning (i.e. conditioned vs unconditioned wasps) on fitness, by measuring the longevity and the fecundity of wasps. Both the nutritional condition and conditioning had significant effects on longevity (Figure 3A; Cox model; nutritional condition: p < 0.001; conditioning: p < 0.001; nutritional condition x conditioning: p = 0.504). Honey diet reduced the risk of death by 16 compared to no food (HR = 0.06), while conditioning increased this risk by 3 (HR = 2.97) indicating a cost of learning and memory formation. The nutritional condition also influenced the fecundity of wasps, so that honey diet increased by 2.75 the number of offspring per female in comparison to no food (Figure 3B; Poisson GLM; diet: p <0.001; conditioning: p = 0.184; diet x conditioning: p = 0.081). Thus overall, early nutritional experience had long lasting effects on adult fitness.

## Discussion

Recent studies showed that a lack of specific nutrients [21, 22] or an unbalanced ratio of these nutrients [20] in food can result in impaired cognitive abilities in model insect species, such as honey bees and fruit flies. Here, we found that olfactory learning performances and memory retention times of *V. canescens* wasps were considerably affected by a poor diet soon after adult emergence. This impact of early adult nutrition on cognition may be particularly critical in this parasitoid species, where females must learn to locate best quality hosts for their progeny.

*V. canescens* can learn a variety of olfactory and visual stimuli associated to their hosts [25, 33, 34]. Here we found that poor diet (sucrose solutions, or no food) at early stages of adulthood significantly reduced olfactory learning and memory retention times. These effects are not developmental as wasps were exposed to nutritional treatments during 24h as adults only. Impaired cognition therefore likely reflects physiological needs for cognitive function in fully developed brains. The fact that conditioning affected longevity, irrespective of the diet, clearly shows the physiological cost associated to learning and memory formation, as previously shown in fruit flies [9].

Unsurprisingly, wasps exposed to food (either honey or sucrose solution) always performed better than starved wasps that presumably lacked energy for basal brain functions. But how can we explain differences in wasps fed honey or sucrose solutions? The fact that wasps exposed to different diets did not differ in their proportions of no choices, indicates that food composition did not affect the locomotion or the motivation of the wasps. Therefore, the observed effects are specific to learning and memory. We discuss two, non-mutually exclusive, possible mechanisms.

Firstly, honey could yield higher cognitive performance because it contains more total nutrients and/or energy. Overall, honey contained much more total energy (284cal/mg) than 20% sucrose solution (80cal/mg) and 10% sucrose solution (40cal/mg). It contained monosaccharaides, like glucose and fructose, that can directly or indirectly (when in combined forms such as glycogen) be absorbed by the insects [35]. These monosaccharaides constitute the carbohydrate-based energy source for the insects. By contrast, sucrose is a disaccharide consisting of one glucose and one fructose molecule that must be broken down by enzymatic reactions before being used as energy source, thus constituting an additional physiological cost [36]. This difference in total amount of carbohydrates in diets may also explain the important difference in survival by wasps fed honey and wasps fed sucrose solution, as carbohydrates have a well-known positive impact on longevity in insects [29, 30, 37].

Alternately, the effects of early adult nutrition on cognition may be due to the lack of specific nutrients in food. Honey contains a rich diversity of nutrients including amino acids and minerals that were not present in sucrose solutions (e.g. Table 1). The most abundant amino acid in honey is proline [38, 39]. The endogenous neutral amino acid L-proline exhibits a variety of physiological and behavioral actions in the nervous system and in increasing or improving memory retention in vertebrates [40, 41]. In honey bees, a decrease of proline in body led to lower learning ability and memory retention [42]. In our experiments, the lack of proteins in sucrose solution likely explains the reduced reproductive success of wasps fed sucrose compared to wasps fed honey, as these are required nutrients for egg production. Honey also includes several macro and micro-elements minerals such as potassium, magnesium, calcium, iron, phosphorus, sodium [43]. Potassium is the most abundant mineral [43, 44]. In human, potassium uptake increase learning and memory [45]. Honey also contains potassium and sodium. The Na+/K+-pump on postsynaptic receptors plays a critical role in synaptic transmission in the brain and a lack of these elements may induced impaired cognitive functions [46, 47, 48].

Unfortunately, our experimental design does not allow to disentangle these mechanisms. Nonetheless, our results demonstrate the crucial importance of adult feeding on their cognitive abilities, longevity and reproduction. These observations in the lab suggest poor adult nutrition can have dramatic consequences for wasps in natural conditions. The importance of nutrition may in fact be greatly magnified in the wild, where wasps must develop costly learning and memory to identify suitable hosts for oviposition, but also need to locate these hosts using olfactory cues associated to the presence of larvae [49]. Wild *V. canescens* wasps are often found in environments where hosts are highly scattered, for instance in orchards where infested fruits are occupied by no more one or two larvae [34, 50]. It is therefore very likely that wasps must flight long distances in order to parasite, thereby incurring additional energetic costs of movements [51]. In these conditions, feeding of high quality foods, such as honey, may provide considerable advantages to wasps. Future experiments could further explore this critical interaction between diet and cognition, using experimental designs of nutritional ecology based on artificial diets controlling for the amount and concentration of nutrients (e.g. [11]). In recent years these approaches have been very successful to identify the effects of specific nutrients and energy contents on fitness traits in many organisms (e.g. flies: [29, 30]; crickets: [37]; mice: [52]), including in hymenoptera (e.g. honey bees: [53]; bumblebees: [54]), and yield considerable promises for investigations in cognition research.

## Material and Methods

### Insect culture

Wasps (*V. canescens*) and their hosts (flour moth *Ephestia kuehniella*) were cultured and tested in incubators at 25°C with a 16:8 Light: Dark photoperiod and 50 ± 5% relative humidity. The *V. canescens* culture originated from wild caught individuals sampled in 2017 (Saveh, Markazi province, Iran) and maintained at the University of Tehran. Natural populations of *V. canescens* contain both thelytokous (asexual) and arrhenotokous (sexual) individuals [55]. Here we only used thelytokous wasps as they are more dependent on nutritional resources acquired as adults (income resources) for reproduction and survival than arrhenotokous conspecifics [27]. *E. kuehniella* eggs were obtained from a laboratory culture at the Insectary and Quarantine Facility of the University of Tehran. *E. kuehniella* larvae were reared on a standard diet made of 48.5g of wheat flour and 3g of brewer yeast [25].

To obtain experimental individuals, groups of 30 one-day old female wasps were presented ca. 200 5^th^ instar host larvae in a large plastic box (30×20×20 cm) and allowed to lay eggs for 24h [25]. Twenty parasitized host larvae were then kept in smaller boxes (5×5×3 cm) until the emergence of adult wasps (range: 25-30 days). Newly emerged wasps (one day old) were fed a 10% honey solution (v/v) to maintain them alive [25]. These wasps (F0) were then placed in one of four nutritional conditions for 24 hours and let to lay eggs. Their progeny (F1) was raised and maintained 24h after emergence in the same nutritional condition before being tested.

### Nutritional conditions

Newly emerged (one day old) female wasps were isolated in glass boxes (10×5×3 cm) and given *ad libitum* access to either: (1) honey (70% carbohydrates: fructose (38% w/v), glucose (30% w/v), 1% fibers, 1.5% protein, total energy (284cal/mg)) (see details in Table 1), (2) 20% sucrose solution (w/v, total energy(80cal/mg)), (3) 10% sucrose solution (w/v, total energy (40cal/mg)), (4) or no food. Wasps were provided honey as droplets on wax-coated strips of paper. Sucrose solutions were provided in gravity feeders (i.e. 4 cm^3^ plastic capsule with a capillary tube inserted at the bottom). Wasps were kept in these boxes for 24h before the behavioral assays.

### Behavioral assays

We performed the cognitive tests in a flight tunnel (200 x 50 x 50 cm) made of transparent Plexiglas (Figure 1; for more details see [56]). The experimental room was illuminated with 2000 lux lights provided by LED lights (Pars Shahab Lamp Co., Iran) [25]. Air was driven through the flight tunnel by a fan located at the upwind end, and extracted outside by a fume hood at the downwind end (wind speed of 70 cm/s). The end opposite to the start zone of the tunnel was divided by a glass separator wall in two decision chambers. Each decision chamber contained an odorant stimulus presented on a filter paper attached to a glass pipette placed vertically on a stand. The behavioral data were recorded though visual observation by an experimenter blind regarding to the nutritional conditions of the wasps.

#### Innate odor preference

To control for any effect of the nutritional condition on odor preference, we assessed the innate odor preference of the wasps. Wasps from each nutritional condition were given a simultaneous choice between two synthetic odors in the flight tunnel: orange and vanilla (97% pure odors: Adonis Gol Darou Group, Iran) [57]. We assumed that our wasp population has never been exposed to these odors prior to the tests, neither in the field nor in the lab. Each odor was presented on a filter paper scented with 1μl of the solution in one of the decision chambers of the tunnel. The wasp was placed at the start zone of the tunnel and allowed to make a choice between the two decision chambers for 15 minutes. Any wasp that spent more than three consecutive minutes within 3 cm around the scented filter paper (landed, walking or hovering around) was considered as “making a choice”. Previous studies show that a wasp landing on an odor site for more than three minutes remains longer than 15 minutes on that site [25]. Any wasp that did not fly in the tunnel within five minutes after the beginning of the test was considered as “making no choice” [25].Fifty wasps were tested for each nutritional condition (N=200 wasps in total). Because we found no innate attraction for either odors, we arbitrarily selected the orange odor as the conditioned stimulus (CS+) and the vanilla odor as the new odor (NOd) in all subsequent experiments.

#### Learning

We assessed the effect of diet on learning performances using olfactory conditioning. To make sure the wasps had some oviposition experience, and thus avoid the inter-individual variability in the sequence and duration of behavioral events associated with learning from the first host encountered [58], female wasps were individually exposed to 15 host larvae (5^th^ instar) for 15 min in a vial (2 cm x 10 cm) before conditioning. Sixty of these wasps were then transferred into conditioning tanks (25 cm x 25 cm x 25 cm) with another 30 host larvae (5^th^ instar). The orange odor (CS+) was pumped into the tanks at an air speed of 1 m/s. The wasps were maintained in these conditions for 2h during which they could associate the orange odor to the presence of host larvae.

Learning performance was assessed 15 min after conditioning by presenting the odors of orange (CS) and vanilla (NOd) in each decision chamber of the flight tunnel. Every wasp that spent more than three consecutive minutes within 3 cm of the CS was considered as making a “correct choice”. Wasps that spent more than three minutes within 3 cm of the NOd made an “incorrect choice”. Wasps that did not fly within five minutes after the beginning of the test made “no choice”. Fifty wasps were tested for each nutritional condition (N=200 wasps in total).

#### Memory retention

We tested the effect of nutritional condition on memory retention time by observing the responses of the conditioned wasps either 2h, 4h, 6h, 8h, 10h, 12h, 14h, 16h, 18h, 20h, 24h, or 30h after conditioning [59]. The responses of the wasps to the CS and the NOd were recorded in the flight tunnel as previously described (see section learning). Fifty wasps were observed in each of the four nutritional conditions and twelve time intervals (N=2400 wasps in total).

### Longevity and fecundity

We tested the effect of the nutritional condition and conditioning on fitness by measuring the longevity and fecundity of conditioned and unconditioned wasps in the four nutritional conditions. To study longevity, we maintained the wasps individually on one of the four nutritional conditions in a plastic box (30 x 20 x 20 cm). We recorded the number of dead wasps every day until all wasps died (18 days). To study fecundity, we placed each wasp in an oviposition cage with 30 host larvae (5^th^ instar). Every day, we monitored the number of wasps emerging from the parasitized hosts and replaced the host larvae by new ones. Thirty females were used for each combination of conditioning and nutritional condition for longevity and for fecundity (N=480 wasps in total).

### Statistical analyses

We analyzed the innate odor responses and learning data using SAS (SAS Institute Inc. 2003). We compared the innate odor response of wasps exposed to different nutritional conditions using Chi-square tests. We tested the effect of nutritional conditions on memory performance using a Generalized Linear Model (GLM) implemented in the procedure GENMOD (binomial family error, logit link function). We compared the least square estimates of the proportions in each level using the Chi-square approximation. When we found a significant effect of the treatment, we applied a Bonferroni’s post hoc multiple comparison tests, and evaluated the two-by-two comparisons at the Bonferroni-corrected significance level of *P* = 0.05/k, where k is the number of comparisons.

We estimated the effect of the nutritional condition on memory retention by developing a dynamic and statistical model following Kishani Farahani et al. [59]. Briefly, the estimation of forgetting relies on a series of observations recorded at different times t_1_; t_2_; …t_n_ after conditioning. At each time, a set of *n*_*t*_ subjects was subjected to a choice test with three possible responses: *a*; *b*; and *c*, which correspond respectively to a preference for the orange side, a preference for vanilla side, and to a no choice. The forgetting of conditioning results in a switch from a high level to a lower level of correct responses, a simultaneous switch from a low level to a high level of no choices, and a switch from a very low to a moderate level of incorrect choices. A constraint links the three responses as *n*_*a*_ *+ n*_*b*_ *+ n*_*c*_ *= n*_*t*_ or *n*_*c*_ *= n*_*t*_ *-n*_*a*_ *-n*_*b*_. The course of these three responses over time can be described by two logistic functions written here as probabilities, *p*_*a*_, *p*_*b*_, *p*_*c*_, constrained by *p*_*a*_ *+ p*_*b*_ *+ p*_*c*_ *=* 1:

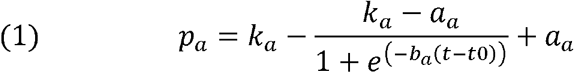

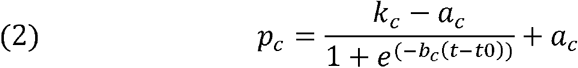

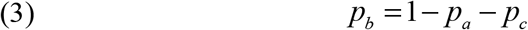

Where *k*_*a*_, respectively *k*_*c*_, and *a*_*a*_, respectively *a*_*c*,_ define the sill and baselines of the logistic models (1) and (2): the baselines are *a*_*a*_ and *a*_*c*_, and the seals are *k*_*a*_ + *a*_*a*_ in model (1), *k*_*c*_ *+ a*_*c*_ in model (2). *k*_*a*_ *+ a*_*a*_ estimates the initial state in model (1), and *a*_*c*_ the final state. It is the inverse in model (2), where a_c_ is the initial state and *k*_*c*_*+a*_*c*_ the final state. A supplementary restriction lies in the fact that, as t0 represents the mean time to oblivion, i.e. the inflection time point of the logistics functions; it has to be the same in all three equations. The data consist of a vector of three counts: *V*_*t*_ =(*n*_*at*_, *n*_*bt*_, *n*_*ct*_) the respective number of subjects responding a; b or c at time t. An R script was written to do this (see Supplementary text S1). The model defined by equations 1 to 3 was fitted individually on each set of ten data. The maximization of the likelihood cannot be fully automatic and requires an initial guess of the seven parameters *k*_*a*_; *a*_*a*_; *b*a; *k*c; *a*_c_; *b*_*c*_; *t0*. This was done by a visual evaluation of each graphic representation of the crossed levels. We compared memory retention times across nutritional conditions using an Analysis of Variance (ANOVA, using SAS).

We analyzed longevity and fecundity data in R 4.0.3 (R Core Team 2020. We tested the effect of the nutritional conditions, conditioning and their interactions on longevity using a Cox proportional hazards regression model (function coxph in package “survival” [60]. We tested the effect of the nutritional conditions, conditioning and their interactions on fecundity using generalized linear mixed-effects model (GLMM) with Poisson family (function glmer in package “lme4” [61]. We added wasp identity and day of experiment as random factors in all models.

## Acknowledgements

This work was funded by Agronutrition CO. (France) and the University of Tehran (Iran). Additionally, SK was supported by a PhD CIFRE grant from Koppert and the French National Agency of Research and Technology (ANRT). ML was supported by the CNRS, two grants from the French National Research Agency (ANR-19-CE37-0024-02 – 3DNaviBee; ANR-20-ERC8-0004-01 – BEEMOVE) and a grant from the European Regional Development Fund (MP0021763 - ECONECT). The sponsors had no involvement in the study design, the collection, and the analysis, the interpretation of the data, the writing, and the decision to submit the manuscript for publication.

## References

1. Healy, S.D., & Jones C.M. Animal learning and memory: An integration of cognition and ecology. Zoology. 105(4), 321–7 (2002).

2. Shettleworth, S. J. Cognition, Evolution, and Behavior (2nd ed). Oxford University Press (2010).

3. Kis, A., Huber, L. & Wilkinson, A. Social learning by imitation in a reptile (Pogona vitticeps). Anim. Cogn. 18, 325–331 (2015).

4. Niven, J. E. & Laughlin, S. B. Energy limitation as a selective pressure on the evolution of sensory systems. J. Exp. Biol. 211, 1792–1804 (2008).

5. Mery, F. & Kawecki TJ. A fitness cost of learning ability in Drosophila melanogaster. Proc. R. Soc. Lond. B. 270(1532), 2465–2469 (2003).

6. Yoshihara, T., Sugihara, K., Kizuka, Y., Oka, S. & Asano, M. Learning/memory impairment and reduced expression of the HNK-1 carbohydrate in beta4-galactosyltransferase-II-deficient mice. J. Biol. Chem. 284(18), 12550–12561 (2009).

7. Bruce K. D., Zsombok A. & Eckel R. H. Lipid Processing in the brain: A key regulator of systemic metabolism. Front. Endocrinol. 8, 60; 10.3389/fendo.2017.00060 (2017).

8. Burns, J.G., Foucaud, J. & Mery, F. Costs of memory: lessons from ‘mini’ brains. Proc. R. Soc. Lond. B. 278, 923–929 (2011).

9. Mery F. & Kawecki TJ. A cost of long-term memory in Drosophila. Science. 308(5725), 1148 (2005).

10. Sutton, M.A. & Schuman, E.M. Dendritic protein synthesis, synaptic plasticity, and memory. Cell. 127(1), 49–58 (2006).

11. Raubenheimer, D. & Simpson, S. Eat Like the Animals: What Nature Teaches Us about the Science of Healthy Eating. Sydney: Harper Collins (2020).

12. Scriba, M. F. et al.. The effect of food quality during growth on spatial memory consolidation in adult pigeons. J. Exp. Biol. 220, 573–581 (2017).

13. Liang, Q. et al.. Postnatal vitamin D intake modulates hippocampal learning and memory in adult mice. Front. Neurosci. 12, 141; 10.3389/fnins.2018.00141 (2018).

14. May, K.A. & Laflamme, D.P. Nutrition and the aging brain of dogs and cat. J. Am. Vet. Med. Assoc. 255(11), 1245–1254 (2019).

15. Beilharz, J.E., Maniam, J. & Morris, M.J. Diet-induced cognitive deficits: The role of fat and sugar, potential mechanisms and nutritional interventions. Nutrients. 7(8), 6719–6738 (2015).

16. 16.Spencer, S.J., Korosi, A., Layé, S., Shukitt-Hale, B., & Barrientos, R.M., Food for thought: How nutrition impacts cognition and emotion. Npj. Sci. Food. 1, 7; 10.1038/s41538-017-0008-y (2017).

17. Giurfa, M. Cognition with few neurons: higher-order learning in insects. Trends Neurosci. 36(5), 285–94 (2013).

18. Perry, C.J., Barron, A.B. & Chittka L. The frontiers of insect cognition. Current Opinion in Behav. Sci. 16, 111–118 (2017).

19. Klein, S., Cabirol, A., Devaud, J.M., Barron, A.B. & Lihoreau, M. Why bees are so vulnerable to environmental stressors. Trends Ecol. Evol. 32(4), 268–278 (2017).

20. Lihoreau, M., Poissonnier, L.A., Isabel, G. & Dussutour, A. 2016. Drosophila females trade off good nutrition with high-quality oviposition sites when choosing foods. J. Exp. Biol. 219, 2514- 2524 (2016).

21. Arien, Y., Dag, A., Zarchin, S., Masci, T. & Shafi, S. Omega-3 deficiency impairs honey bee learning. PNAS. 112 (51), 15761–15766 (2015).

22. Arien Y, Dag A, Shafir S. Omega-6:3 ratio more than Aasolute lipid level in diet affects associative learning in honey bees. Front. Psychol. 9, 1001; 10.3389/fpsyg.2018.01001 (2018).

23. Thiel, A., Schlake, S. & Kosior, D. Omnia tempus habent: habitat-specific differences in olfactory learning and decision making in parasitic wasps. Anim. Cogn. 16(2), 223–32 (2003).

24. Froissart, L., Giurfa, M., Sauzet, S. & Desouhant, E. Cognitive adaptation in asexual and sexual wasps living in contrasted environments. PLOS ONE. 12(5), e0177581. 10.1371/journal.pone.0177581 (2017).

25. Kishani Farahani, H., Moghadassi, Y., Alford, L. & van Baaren, J. Effect of interference and exploitative competition on associative learning by a parasitoid wasp: a mechanism for ideal free distribution? Anim. Behav. 151, 157–163 (2019).

26. Desouhant, E., Driessen, G., Amat, I. & Bernstein, C. Host and food searching in a parasitic wasp Venturia canescens: a trade-off between current and future reproduction? Anim. Behav. 70 (1), 145–152 (2005).

27. Pelosse, P., Bernstein, C. & Desouhant, E. Differential energy allocation as an adaptation to different habitats in the parasitic wasp Venturia canescens. Evol. Ecol. 21, 669–685 (2007).

28. Benelli, G., Giunti, G., Tena, A. Desneux, N., Caselli, A. & Canale, A. The impact of adult diet on parasitoid reproductive performance. J. Pest. Sci. 90, 807–823 (2017).

29. Lee, J.C. & Heimpel, G.E. Floral resources impact longevity and oviposition rate of a parasitoid in the field. J. Anim. Ecol. 77, 565–572 (2008a).

30. Lee, J.C. & Heimpel, G.E. Effect of floral nectar, water, and feeding frequency on Cotesia glomerata longevity. BioControl. 53, 289–294 (2008b).

31. Rusch, A., Suchail, S., Valantin-Morison, M., Sarthou, J.P. & Roger-Estrade, J. 2013. Nutritional state of the pollen beetle parasitoid Tersilochus heterocerus foraging in the field. BioControl. 58 (1), 17-26 (2013).

32. Campos, M. G. R. et al.. Pollen composition and standardization of analytical methods. J. Apic. Res. 47(2), 156–163 (2008).

33. Lucchetta, P., Bernstein, C., Théry, M. Lazzari, C. & Desouhant, E. Foraging and associative learning of visual signals in a parasitic wasp. Anim. Cogn. 11, 525–533 (2008).

34. Thiel, A., Schlake, S. & Kosior, D. Omnia tempus habent: habitat-specific differences in olfactory learning and decision making in parasitic wasps. Anim. Cogn. 16, 223–232 (2013).

35. Rittschof, C.C. & Schirmeier S. insect models of central nervous system energy metabolism and its links to behavior. Glia. 66(6), 1160–1175 (2017).

36. Sammataro, S., Weiss, M. 2013. Comparison of productivity of colonies of honey bees, Apis mellifera, supplemented with sucrose or high fructose corn syrup, J. Insect Sci. 13(1), 19; 10.1673/031.013.1901 (2013).

37. Maklakov, A.A. et al.. Sex-specific fitness effects of nutrient intake on reproduction and lifespan. Curr. Biol. 18(14), 1062–1066 (2008).

38. Di Girolamo, F., D’Amato, A. & Righetti, P.G. Assessment of the floral origin of honey via proteomic tools. J. Proteom. 75(12), 3688–93 (2012).

39. Kečkeš, J. et al.. Amino acids profile of Serbian unifloral honeys. J. Sci. Food Agric. 93, 3368–3376 (2013).

40. Wyse AT, Netto CA. Behavioral and neurochemical effects of proline. Metab. Brain. Dis. 26(3), 159–172 (2011).

41. Schulz, D., Morschel, J., Schuster, S., Eulenburg, V. & Gomeza, J. Inactivation of the mouse L-proline transporter PROT alters glutamatergic synapse biochemistry and perturbs behaviors required to respond to environmental changes. Front. Mol. Neurosci. 11, 279; 10.3389/fnmol.2018.00279 (2018).

42. Gage S. L. et al.. Nosema ceranae parasitism impacts olfactory learning and memory and neurochemistry in honey bees (Apis mellifera). J. Exp. Biol. 19, 221; 10.1242/jeb.161489 (2018).

43. Solayman, M. et al.. Physicochemical properties, minerals, trace elements, and heavy metals in honey of different origins: A comprehensive review. Compr. Rev. Food. Sci. Food. Saf. 15, 219–233 (2016).

44. Yücel, Y. & Sultanoğlu, P. 2013. Characterization of Hatay honeys according to their multi-element analysis using ICP-OES combined with chemometrics. Food Chem. 140(1-2), 231-237 (2013).

45. Cisternas, P. et al.. 2015. The increased potassium intake improves cognitive performance and attenuates histopathological markers in a model of Alzheimer’s disease, BBA-Mol. Basis Dis. 1852(12), 2630-2644 (2015).

46. Pivovarov, A.S., Calahorro, F. & Walker, R.J. Na+/K+-pump and neurotransmitter membrane receptors. Invert. Neurosci. 19(1), 1; 10.1007/s10158-018-0221-7 (2018).

47. Gardoni, F. et al.. Decreased NR2B subunit synaptic levels cause impaired long-term potentiation but not long-term depression. J. Neurosci. 29(3), 669–77 (2009).

48. Zhang, L. et al.. Na(+)/K(+)-ATPase inhibition upregulates NMDA-evoked currents in rat hippocampal CA1 pyramidal neurons. Fundam. Clin. Pharmacol. 26(4), 503–12 (2012).

49. Vet, L. E. M., Wackers, F. L. & Dicke, M. How to hunt for hiding hosts - the reliability-detectability problem in foraging parasitoids. Netherlands J. Zool. 41, 202–213 (1991).

50. Driessen, G. & Bernstein, C. Patch departure mechanisms and optimal host exploitation in an insect parasitoid. J. Anim. Ecol. 68(3), 445–59 (1999).

51. Amat I. et al.. Fuelling flight in a parasitic wasp: which energetic substrate to use? Ecol. Entomol. 37(6), 480–489 (2012).

52. Solon-Biet, S.M. et al.. 2015. Macronutrients, reproduction, and lifespan in mice. PNAS. 112(11), 3481-3486 (2015).

53. Altaye, S.Z., Pirk, C.W., Crewe, R.M. & Nicolson, S.W. Convergence of carbohydrate-biased intake targets in caged worker honeybees fed different protein sources. J. Exp. Biol. 213(19), 3311–3318 (2010).

54. Kraus, S. et al.. Bumblebees adjust protein and lipid collection rules to the presence of brood. Curr. Zool. 65, 437–446 (2019).

55. Foray, V., Henri, H., Martinez, S., Gibert, P. & Desouhant, E.. Occurrence of arrhenotoky and thelytoky in a parasitic wasp Venturia canescens (Hymenoptera: Ichneumonidae): Effect of endosymbionts or existence of two distinct reproductive modes? Eur. J. Entomol. 110(1), 103–107 (2013).

56. Drost, Y.C., Lewis, W.J. & Tumlinson, J.H. Beneficial arthropod behavior mediated by airborne semiochemicals. J. Chem. Ecol. 14, 1607–1616 (1988).

57. Scotto-Lomassese, S. et al.. Suppression of adult neurogenesis impairs olfactory learning and memory in an adult insect. J. neurosci. 23(28), 9289–9296 (2003).

58. Mills, N.J. & Kuhlmann, U. Oviposition behavior of Trichogramma platneri Nagarkatti and Trichogramma pretiosum Riley (Hymenoptera: Trichogrammatidae) in patches of single and clustered host eggs. Biol. Control. 30, 42–51 (2004).

59. Kishani Farahani, H. et al. Early life nutritional quality effects on adult memory retention in a parasitic wasp. Behav. Ecol. 28(3), 818–826 (2017).

60. Therneau, T. A Package for Survival Analysis in R. R package version 3.2-7, https://CRAN.R-project.org/package=survival (2020).

61. Bates, D., Maechler, M., Bolker, B. & Walker, S. Fitting Linear Mixed-Effects Models Using lme4. J. Stat. Softw. 67(1), 1–48 (2015).

